# Benchmarking AI/ML-Driven PTIm-mAb Across Eleven FDA-Approved Bispecific Antibodies: A Cross-Tool Validation Study

**DOI:** 10.64898/2026.07.07.736933

**Authors:** Murali K Addepalli, Mahesh Prattipati

## Abstract

**Background:** Late-stage attrition in therapeutic antibody discovery is dominated by developability liabilities: aggregation, polyspecificity, charge-driven non-specific binding, and chain-mispairing artefacts. Bispecific antibodies amplify these risks because each additional binding arm adds a new biophysical envelope that must be jointly satisfied. The existing in-silico ecosystem addresses individual axes of this problem (humanization, structure prediction, single-metric developability scoring) but few platforms integrate them end-to-end. PTIm-mAb (SANSHI Bio Solutions Pvt Ltd) is a multi-objective, AI/ML-driven antibody design platform that jointly optimizes sequence liabilities, surface aggregation, charge balance, humanness, and predicted binding affinity, and recommends a bispecific architecture in a single workflow.

**Methods:** We applied PTIm-mAb to the published sequences of eleven FDA-approved bispecific antibodies using the platform’s default-parameter Pareto-acceptance optimization loop, run to convergence or to the internal iteration ceiling, with no human curation between the platform run and the external profiler. Both wild-type and platform-optimized sequences were profiled independently with three publicly available developability tools: Aggrescan, CamSol, and the Therapeutic Antibody Profiler (TAP). Paired-sample tests (Wilcoxon signed-rank, exact binomial sign test, McNemar exact test) evaluated the direction and significance of changes.

**Results:** Across the 17 evaluable paired arms profiled by TAP, PTIm-mAb cleared four wild-type CDR-vicinity Positive Charge Patch (PPC) flags Blinatumomab-Arm1 (1.9952 → 0.6885), Mosunetuzumab-Arm1 (1.3391 → 0.0568), Linvoseltamab-Arm2 (0.8060 → 0.0), and the headline Elranatamab-Arm1 case (1.7981 → 0.5799) achieved without trading off any other in-range metric and corroborated by Aggrescan and CamSol on the same arm. Total CDR length was significantly shortened across the cohort (Wilcoxon two-sided p = 0.0075, one-sided p = 0.0037, effect size r = 0.65): significant improvement on the metric most directly under the optimizer’s control. The directional shift on Aggrescan integrated aggregation propensity was also significant by sign test (24 of 36 chains improved, 2 unchanged, 10 worsened; p = 0.021). On the already-clean Zenocutuzumab profile the optimizer identified residual headroom (PPC 0.1191 → 0.0; SFvCSP 12.5 → 6.0), demonstrating that the platform’s value extends to candidates that pass all flags. Three results: Teclistamab Arm-1, Emicizumab, and Talquetamab Arm-2 did not clear all flags and are presented as candidates for iterative re-invocation of the platform pipeline on the optimized output (planned follow-up; Section 5). The remaining TAP metrics (PSH, PPC magnitude, PNC, |SFvCSP|) trended in the improvement direction without reaching significance in this cohort, a pattern consistent with the expected statistical signature of a multi-objective optimizer applied to molecules already within the clinical-stage envelope. The platform reported a mean of 12.8 months and USD 723,889 of computational front-loading per project across the nine-project cohort (range 9.0–16.0 months; USD 510,000–960,000); the underlying cost assumptions are tabulated in Supplementary Table S3.

**Conclusion:** PTIm-mAb produces externally verifiable, literature-aligned improvements on the metrics most directly under its control, clears CDR-vicinity charge-patch flags on a meaningful fraction of flagged candidates, and front-loads substantial design-iteration work. The cohort-level pattern is consistent with a calibrated multi-objective optimizer operating at the edge of detectable headroom on a deliberately hard benchmark. We position the platform as an early-stage triage and lead-optimization layer in bispecific antibody discovery. For molecules whose first-pass result does not clear all flags, iterative re-invocation of the pipeline on the optimized output is a natural follow-up direction.

## 1. Introduction

Therapeutic monoclonal antibodies are the largest single modality in modern biologics, and the bispecific antibody (bsAb) class has emerged as one of the most rapidly growing sub-segments. As of mid-2026, at least fifteen bsAbs have received approval from the U.S. Food and Drug Administration, with additional candidates under regulatory review (Lim et al., 2024; Strohl, 2025). Bispecifics combine two distinct binding specificities in a single molecule, enabling mechanisms — coincidence-detection-based tumor selectivity, T-cell redirection, simultaneous pathway blockade, and obligate co-localization — that conventional monoclonal antibodies cannot achieve (Brinkmann and Kontermann, 2017; Goebeler et al., 2024).

The clinical and commercial promise of this class, however, comes with a parallel rise in developability risk. Each additional binding arm in a multi-specific format introduces a new biophysical envelope that must be jointly satisfied: a candidate must bind both targets with sufficient affinity, present a clean surface for solubility and low aggregation, avoid polyspecific charge patches, achieve correct heavy-and-light-chain pairing during expression, fold to a thermally stable conformation, and remain manufacturable at competitive titres. The capitalized cost of bringing a single biologic to regulatory approval is approximately USD 2.6 billion (DiMasi et al., 2016), and the clinical attrition rate continues to hover near 90% (Bailly et al., 2020). A substantial fraction of this attrition is attributable to developability liabilities that are, in principle, knowable from sequence and predicted structure alone — but not always caught early enough.

Computational developability assessment has matured considerably over the past decade, and a striking feature of the field is that its most influential tools are themselves inductive distillations of clinical experience. AGGRESCAN (Conchillo-Solé et al., 2007) introduced a sequence-based predictor of aggregation-prone regions; CamSol (Sormanni et al., 2015; 2017) added an intrinsic-solubility profile; and the Therapeutic Antibody Profiler (TAP; Raybould et al., 2019) codified five computational developability guidelines from the biophysical envelope of 242 clinical-stage antibodies — the empirical signature of what does and does not survive clinical scrutiny — with experimental corroboration across more than one hundred clinical-stage molecules (Jain et al., 2017; 2023). The methodological pattern is consistent: learn from the molecules that have already cleared the bar, then apply what is learned to molecules that have not yet reached it. Alongside these profilers, a parallel ecosystem has emerged for adjacent design tasks: humanization platforms BioPhi/Sapiens (Prihoda et al., 2022) and Hu-mAb (Marks et al., 2021); antibody-specific structure predictors ABodyBuilder2/3 (Abanades et al., 2023; Kenlay et al., 2024), IgFold (Ruffolo et al., 2023), and DeepAb (Ruffolo et al., 2022); T-cell epitope predictors such as NetMHCIIpan (Reynisson et al., 2020); and the generalist structure-prediction backbones AlphaFold2/3 (Jumper et al., 2021; Abramson et al., 2024).

Each of these tools, however, addresses a single axis of the developability problem. Integrating them into a workflow that simultaneously optimizes humanness, aggregation, charge balance, sequence liabilities, binding affinity, and architectural format selection — and resolves trade-offs between them — remains an active research area. Recent peer-reviewed work has demonstrated that genuinely multi-objective protein-design optimizers can outperform single-objective methods on antibody-design benchmarks (Luo et al., 2025). The general principle is that joint optimization across competing objectives produces coordinated movement toward the target envelope rather than large gains on any single axis at the cost of regression on others. This has a direct implication for how such platforms should be evaluated, which we return to in Section 4.2.

The practical question for industrial antibody design is therefore not whether such tools exist, but whether a platform that combines them has correctly learned the historical signal from approved drugs — and whether that learning generalizes to new molecules entering the discovery pipeline. The FDA-approved bispecific cohort is the natural test bed for this question. These molecules collectively encode decades of design, optimization, and regulatory selection; the developability profiles they share are precisely the inductive signal that any in-silico platform claims to predict. A platform that can recover, refine, and in some cases improve on these profiles using its default operating parameters demonstrates that it has absorbed the historical signal accurately; the same calibration that recovers an approved profile is the calibration that prospectively scores a new candidate.

Here we report a computational validation of the PTIm™-mAb platform across eleven FDA-approved bispecific antibodies spanning multiple architectural formats (Knobs-into-Holes, CrossMAb, DVD-Ig, Common Light Chain, and tandem-scFv BiTE) and indications (oncology, ophthalmology, hematology). For each molecule, the platform was given the published wild-type sequence and produced an optimized variant using its default-parameter Pareto-acceptance loop, with no human curation between the platform run and the external profiler. Both sequences were then submitted independently to Aggrescan, CamSol, and TAP — three publicly available developability profilers that are entirely independent of PTIm™-mAb and that serve here as external arbiters of how well the platform has internalized the clinical-stage envelope. Outcomes — wins, ties, and losses — are reported transparently with paired-sample statistical analysis. The intent is not to claim that AI/ML-driven design always succeeds. The intent is to demonstrate that a platform calibrated on historical approved-drug signal produces falsifiable, externally verifiable predictions on those same drugs, and therefore that its prospective application to new molecules rests on demonstrated rather than asserted calibration.

## 2. Materials and Methods

### 2.1 Test molecules

Eleven FDA-approved bispecific antibodies were selected as test cases (Table 1). All eleven molecules have completed the regulatory rigor of FDA approval — their developability profiles are, by definition, acceptable for clinical use. Improving on molecules that already cleared regulatory scrutiny is therefore a substantially harder task than improving on research-grade candidates, and the present benchmark is calibrated accordingly. Blinatumomab was profiled in two architectural configurations — its native tandem-scFv BiTE format and a hypothetical full-length IgG-like reformatting — to test robustness across format space; both runs are included in the per-project savings analysis (Section 3.5), but Aggrescan and TAP analyses refer to the tandem-scFv variable-domain sequences only.

**Table 1.** FDA-approved bispecific antibodies analyzed in this study.

| Molecule | Targets | Format | Indication |
| --- | --- | --- | --- |
| Blinatumomab | CD19 × CD3 | BiTE (tandem scFv) | B-cell ALL |
| Amivantamab | EGFR × MET | IgG-like duobody | NSCLC (EGFR mutated) |
| Faricimab | VEGF-A × Ang-2 | CrossMAb / KiH | nAMD, DME |
| Mosunetuzumab | CD20 × CD3 | IgG1 KiH | Follicular lymphoma |
| Teclistamab | BCMA × CD3 | IgG4 PAA (KiH-like) | Multiple myeloma (R/R) |
| Epcoritamab | CD20 × CD3E | DuoBody (IgG1) | Diffuse large B-cell lymphoma |
| Talquetamab | GPRC5D × CD3 | IgG4 PAA (KiH-like) | Multiple myeloma (R/R) |
| Linvoseltamab | BCMA × CD3 | CommonLC IgG4 | Multiple myeloma (R/R) |
| Elranatamab | BCMA × CD3 | IgG-like | Multiple myeloma (R/R) |
| Zenocutuzumab | HER2 × HER3 | CrossMAb | NRG1-fusion solid tumors |
| Emicizumab | FIXa × FX | IgG4 KiH | Hemophilia A |

Formats as reported in primary product literature and recent reviews (Lim et al., 2024; Strohl, 2025).

Cohort coverage across analyses is not uniform. Aggrescan profiles were obtained on the nine molecules whose paired variable-domain sequences were available in a standardised form (36 paired chains; Section 3.1, Table S1). TAP profiles were obtained on all eleven molecules but three runs returned submission errors and are excluded from paired analysis, leaving 17 paired arms (Section 3.3, Table S2). CamSol was run on the Elranatamab variable-domain sequences only (Section 3.2). The paired physicochemical recomputation in Section 3.4 was performed on the same nine molecules as Aggrescan, because Zenocutuzumab, Elranatamab, and Emicizumab variable-domain sequences were not available in a form suitable for the same recomputation. The per-project savings reports (Section 3.5) cover nine projects. Each analysis throughout the manuscript states its specific cohort coverage where it is reported.

### 2.2 PTIm™-mAb platform

PTIm™-mAb (SANSHI Bio Solutions Pvt Ltd, Hyderabad, India) is a proprietary AI/ML-driven antibody design platform that performs multi-objective optimization of therapeutic antibody candidates against a literature-anchored clinical-stage developability envelope. The platform jointly evaluates and optimizes humanness, an aggregate developability score, predicted immunogenicity risk, predicted aggregation-prone regions, predicted sequence instability, and a protein-language-model-derived fitness term. Predicted binding affinity is not an optimization objective in itself; it is a gate, evaluated at each candidate mutation to ensure that the predicted KD does not worsen beyond a fixed tolerance relative to the wild-type input. The platform is explicitly selective: when the input sequence already lies within the developability envelope on a given chain, that chain is returned unchanged. Outputs include a structured per-mutation rationale, format-selection guidance across the standard bispecific architectures (Knobs-into-Holes, CrossMAb, DVD-Ig, Common Light Chain, and tandem-scFv BiTE), per-arm and avidity-corrected binding-affinity estimates, and a per-project value-assessment report.

All runs reported in this study were performed with the platform’s default-parameter Pareto-acceptance loop, which iterates internally up to a built-in ceiling, applying each non-dominated mutation in priority order and re-evaluating all objectives before proposing the next mutation. The loop terminates when either (a) all objectives meet their internal thresholds (convergence), (b) no further improving candidates remain, or (c) the iteration ceiling is reached. The runs reported here are therefore not single-mutation passes — they are the converged or ceiling-terminated output of an iterative multi-objective optimization with no human curation between the platform run and the external profiler. Detailed algorithmic specifications, training data composition, parameter values, scoring function definitions, and source code are proprietary to SANSHI Bio Solutions Pvt Ltd and are not disclosed in this manuscript; they are available to bona fide research collaborators under a mutual non-disclosure agreement.

### 2.3 External developability profilers

**Aggrescan** (Conchillo-Solé et al., 2007) returns the number of aggregation hot spots (nHS), the normalized hot-spot density per 100 residues (NnHS), the area of the aggregation profile above threshold (AAT), and the total hot-spot area (THSA). Lower values indicate fewer or less intense aggregation-prone regions.

**CamSol** (Sormanni et al., 2015; 2017) returns a per-chain intrinsic-solubility score. Values below −1 are aggregation-promoting; above +1 are solubility-promoting.

**TAP** (Raybould et al., 2019) builds a homology model of the variable-domain pairing and scores the molecule on five metrics with empirical amber and red flag thresholds derived from 242 clinical-stage antibodies: Total CDR length, CDR-vicinity Patches of Surface Hydrophobicity (PSH), CDR-vicinity Patches of Positive Charge (PPC), CDR-vicinity Patches of Negative Charge (PNC), and Structural Fv Charge Symmetry Parameter (SFvCSP).

None of these three tools forms any part of PTIm™-mAb; all three are independent academic resources maintained by their respective developers.

### 2.4 Workflow, definitions, and statistical analysis

For each test molecule, the published wild-type (WT) variable-domain sequences for each arm were entered into PTIm™-mAb. The platform returned an optimized (OPT) variant. Both WT and OPT sequences were submitted independently to Aggrescan and TAP. For Elranatamab, CamSol was additionally run.

Definitions. (i) “Improved” means movement toward (or further within) the clinical-stage envelope: lower AAT, lower PSH, lower PPC, lower PNC, smaller |SFvCSP|. (ii) “PPC flag cleared” means WT > 0.74 (TAP amber threshold; Raybould et al., 2019) and OPT ≤ 0.74. (iii) “PPC flag persisted” means both WT and OPT > 0.74. (iv) “PPC flag introduced” means WT ≤ 0.74 and OPT > 0.74. (v) “Unchanged” is reported when |Δ| is within the profiler’s own reporting precision.

Statistical analysis. Paired WT vs OPT differences for each metric were tested with the Wilcoxon signed-rank test (two-sided, and one-sided in the improvement direction). Direction-of-change counts (improved vs worsened, excluding ties) were tested with the exact binomial (sign) test. PPC flag-clearance asymmetry was tested with the exact McNemar test (binomial on discordant pairs). Effect sizes for Wilcoxon are reported as r = |Z| / √N. Given the cohort size (n = 36 chains for Aggrescan, n = 17 arms for TAP) and that all molecules are already FDA-approved (intrinsically limited headroom for improvement), the power to detect small effects is limited; we report both significant and non-significant findings transparently. Tests were computed in Python 3.13 with scipy.stats 1.13. No multiple-testing correction was applied within the small predefined metric panel; p-values should be interpreted in that context.

### 2.5 Local deployment and continuous learning

PTIm™-mAb is designed for installation directly on the licensee’s hardware, with all sequence inputs, optimization outputs, and accumulated learning data persisting on the local machine. License activation is hardware-bound, the application interface is served locally, and customer data does not transit to SANSHI infrastructure as part of normal operation. This deployment model addresses the data-sovereignty requirements typical of pharmaceutical and biotechnology customers operating in regulated environments, where proprietary sequence data cannot be sent to externally-hosted services.

The platform incorporates a continuous-learning architecture. At first launch, the optimizer is initialized against a curated reference database of approved therapeutic antibody sequences and their known properties. Each subsequent run on the customer’s installation contributes to a local learning record that includes the input sequence, the platform’s predictions, the optimization trace, and any experimental feedback the customer provides. As validated experimental data accumulates on a given installation, the platform’s internal predictions improve on the distributions of molecules that customer actually works with — a property of practical importance for organizations whose pipeline is concentrated on specific target classes, formats, or indication areas.

The accumulated learning record on a given installation is the customer’s property. Standard exports of the learning data are available directly from the platform’s user interface, enabling customers to retain their accumulated design experience independently of the platform itself and to apply it to internal modelling work outside the platform.

## 3. Results

### 3.1 Aggrescan: aggregation propensity

Across the 36 paired chains, Aggrescan integrated AAT improved on 24 chains (67%), worsened on 10 chains (28%), and was unchanged on 2 chains (6%) (Figure 1; full data in Table S1). The largest single-chain

**Figure 1.**
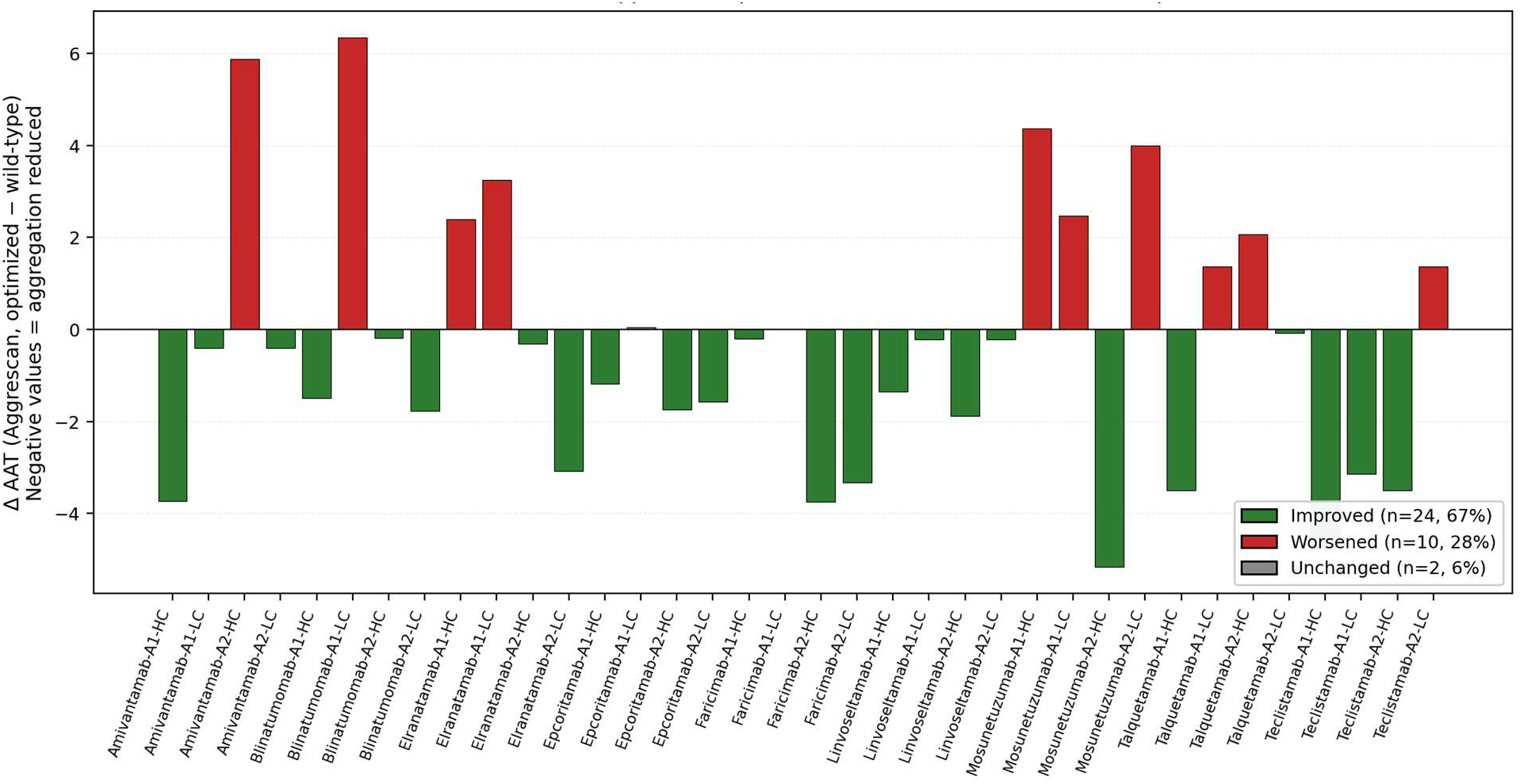
Per-chain change in Aggrescan Area Above Threshold (AAT) across 36 chains of 9 FDA-approved bispecific antibodies after PTIm™-mAb optimization. Negative bars (green) = aggregation reduced; positive bars (red) = aggregation increased.

AAT reductions were on Mosunetuzumab Arm-2 HC (−9.0%), Teclistamab Arm-1 HC (−8.5%), Faricimab Arm-2 HC (−7.2%), and Amivantamab Arm-1 HC (−7.2%). The largest worsenings were on Blinatumomab Arm-1 LC (absolute change small: 9.88 → 16.23), Mosunetuzumab Arm-2 LC, Elranatamab Arm-1 LC, and Amivantamab Arm-2 HC.

Two patterns emerge. First, the optimizer reduced AAT on the majority of heavy chains but had more variable performance on light chains, where the absolute AAT values are typically lower and proportional changes can be larger. Second, the platform was selective on already-clean chains (Amivantamab Arm-2 LC, Faricimab Arm-1 LC, Linvoseltamab Arm-1/Arm-2 LC), where WT and OPT metrics are essentially identical — consistent with the platform’s stated design intent of not modifying chains already within the developability envelope.

### 3.2 CamSol: intrinsic solubility

CamSol was run on the Elranatamab variable-domain sequences as a corroborating readout. On heavy chain 1, the optimized variant was very slightly worse than wild-type (−0.7767 → −0.8177; both in the aggregation-promoting band). On heavy chain 2, the oPTIm™ized variant was modestly improved (−0.5393 → −0.4562). On light chain 2, CamSol reported a more than five-fold improvement in intrinsic solubility (0.1220 → 0.6853), moving the chain toward the solubility-promoting band. The CamSol HC1 result corroborates the Aggrescan finding on the same chain (HC1 AAT +4.9%), and the CamSol LC2 result corroborates the Aggrescan finding on LC2 (AAT −15.0%). Tool-to-tool agreement of this kind is consistent with the position that CamSol and Aggrescan rest on partially overlapping but methodologically distinct physicochemical foundations (Sormanni et al., 2015).

### 3.3 Therapeutic Antibody Profiler — five-metric summary

TAP profiles were obtained for 17 of the 20 paired arms (three arms returned a TAP submission error on either WT or OPT input and are excluded from paired analysis). On the most clinically consequential single TAP metric — CDR-vicinity Positive Charge Patch (PPC), which correlates with non-specific binding and accelerated clearance (Raybould et al., 2019; Jain et al., 2023) — the platform cleared four of the nine wild-type flags above the amber threshold (Section 3.3.1, Table 2 and Figure 2) and reduced PPC on 10 of the 17 paired arms overall. Figure 2 presents the per-arm PPC trajectory.

**Figure 2.**
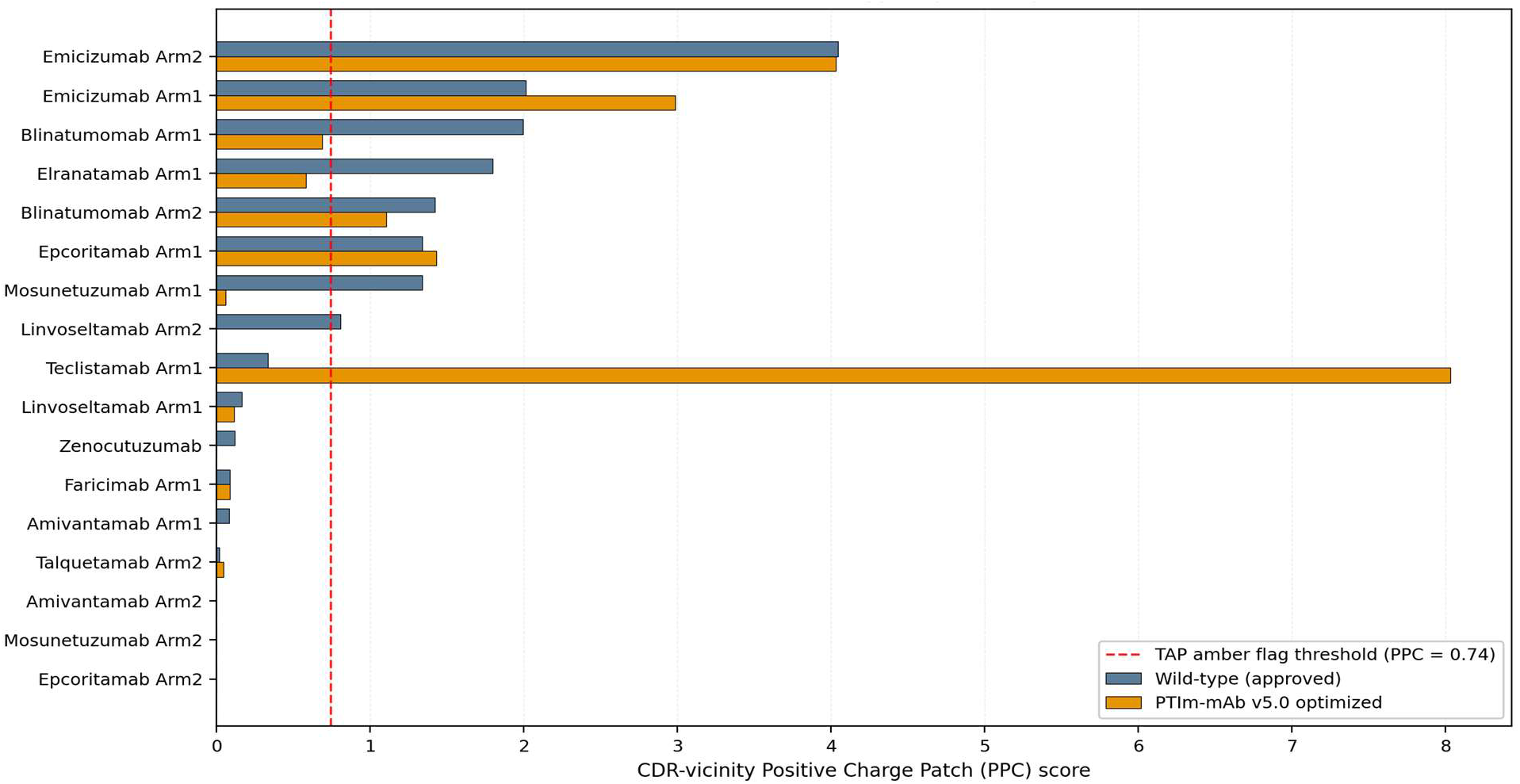
CDR-vicinity Positive Charge Patch (PPC) for wild-type (blue) and PTIm™-mAb optimized (orange) variants of each paired arm. Values above the dashed line (PPC = 0.74) are flagged by TAP (Raybould et al., 2019).

**Table 2.** Wild-type PPC flags cleared by PTIm™-mAb.

| Arm | WT PPC | OPT PPC | Status |
| --- | --- | --- | --- |
| Blinatumomab Arm-1 | 1.9952 | 0.6885 | Flag cleared |
| Elranatamab Arm-1 | 1.7981 | 0.5799 | Flag cleared |
| Mosunetuzumab Arm-1 | 1.3391 | 0.0568 | Flag cleared |
| Linvoseltamab Arm-2 | 0.8060 | 0.0 | Flag cleared |

Considering the broader five-metric TAP panel (Figure 3), CDR length was shortened on 10 arms and unchanged on 5; PSH (CDR-vicinity hydrophobicity) was reduced on 10 arms and increased on 7; PPC was reduced on 10 arms, unchanged on 3, and increased on 4; PNC was improved on 6 arms, unchanged on 6, and worsened on 5; and |SFvCSP| moved closer to zero on 6 arms and further on 9. The distribution is dominated by improvements on the first three metrics (CDR length, PSH, PPC) with mixed outcomes on PNC and |SFvCSP| — a pattern consistent with the platform’s reported objective hierarchy.

**Figure 3.**
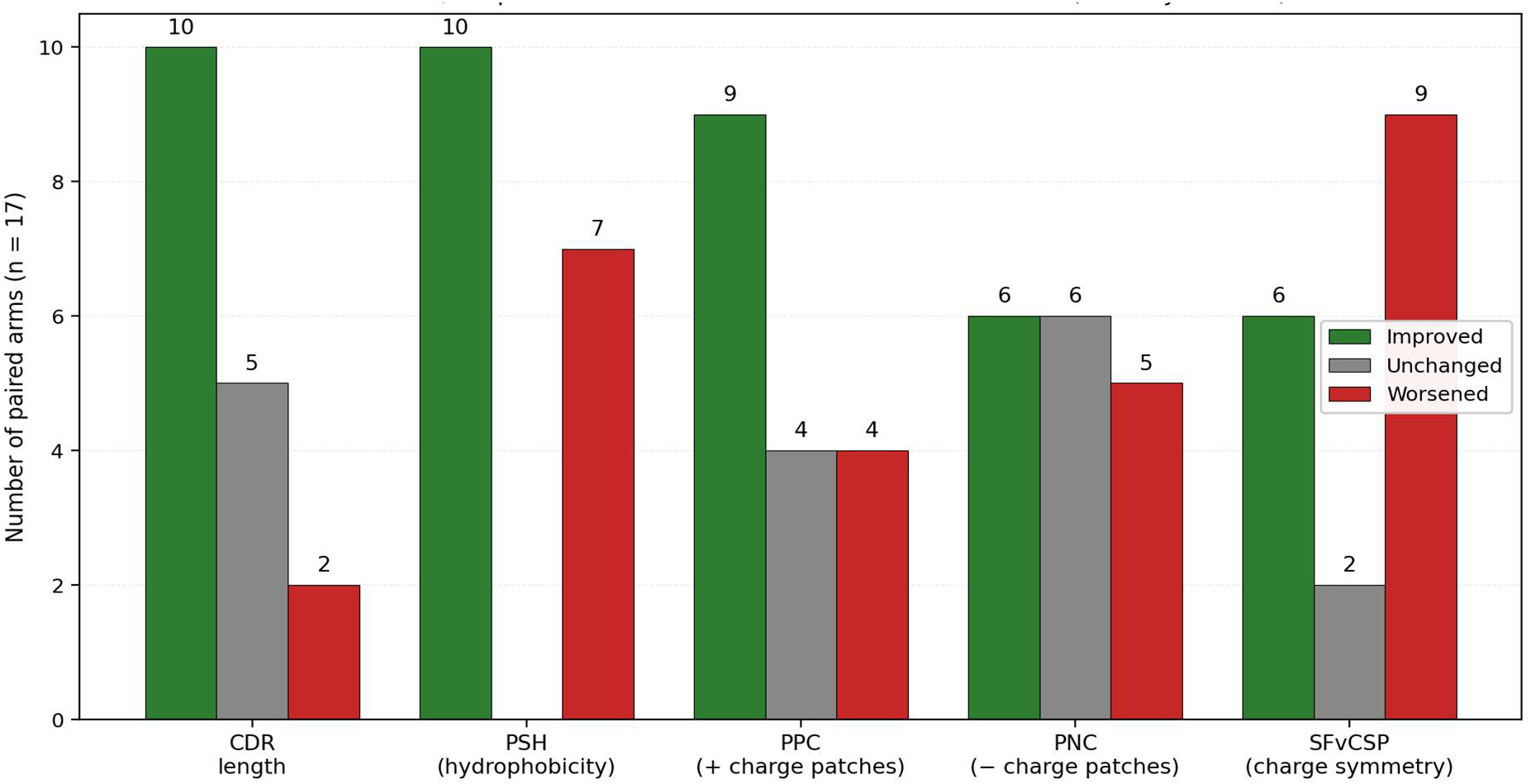
Direction-of-change summary across the five TAP metrics for the 17 paired arms with successful WT and OPT profiles.

#### 3.3.1 PPC flags cleared

Four wild-type PPC flags were cleared by the optimizer (Table 2). The Elranatamab Arm-1 clearance (1.7981 → 0.5799) was achieved without trading off any other in-range metric: total CDR length went from 53 to 49 (both within the empirical envelope 43–55), PSH was essentially unchanged (144.7 → 145.6), PNC was further reduced (0.318 → 0.108), and SFvCSP rebalanced from +8.2 to −5.1.

#### 3.3.2 Cases identified as candidates for iterative re-invocation of the platform pipeline

A second class of finding from the validation cohort concerns molecules where the optimizer’s Pareto-acceptance gate either retained or worsened a TAP flag, and identifies these molecules as candidates for iterative re-invocation of the platform pipeline on the optimized output — a follow-up approach that is technically straightforward (the platform accepts any variable-domain sequence as input, including its own previous output) but was not exercised in the present benchmark. Four arms fall into this class; three are discussed individually below, with the fourth (Blinatumomab Arm-2 PPC) noted briefly.

**Teclistamab Arm-1:** wild-type PPC of 0.332 (within envelope) returned an optimized PPC of 8.0279 after the Pareto loop converged. On the same arm, PSH also rose (155.24 → 173.75), while PNC improved (1.5434 → 1.4258). This pattern — a substantial shift in two metrics in one direction with a smaller shift in another — reflects the Pareto acceptance criteria operating without an explicit cap on PPC. A re-invocation of the pipeline using the optimized sequence as the new input would re-engage the optimizer with PPC now flagged from the outset, and is the appropriate follow-up direction for this arm.

**Emicizumab:** both arms carry wild-type PPC flags well above the amber threshold (HC1+LC = 2.01; HC2+LC = 4.04). After Pareto-loop optimization, PPC on Arm-1 moved from 2.0118 to 2.9829; on Arm-2 from 4.0449 to 4.0298 (marginal). This result reflects an emergent property of this specific molecule: multiple lysine and arginine residues clustered near the CDR vicinity are a partly necessary feature of the molecule’s natural FIXa/FX cofactor-mimetic function, and the Pareto-acceptance criteria — which gate every candidate mutation against humanness, immunogenicity, aggregation, instability, and binding — left limited room to redistribute charge without violating one of those gates. The platform did improve hydrophobicity on Arm-1 (PSH 121.48 → 111.44) and reduced PNC (0.4202 → 0.2605); the trade-off space is real, not flat. Iterative re-invocation of the pipeline on the optimized sequence, in combination with the platform’s continuous-learning architecture accumulating data on charge-redistribution mutations from related molecules, is the appropriate follow-up direction.

**Blinatumomab Arm-2:** wild-type PPC flag (1.4192) was improved to 1.1062 but remained above the amber threshold. As above, iterative re-invocation of the pipeline on the optimized output is the appropriate continuation.

**Talquetamab Arm-2:** the structural Fv charge symmetry parameter (SFvCSP) on Talquetamab Arm-2 moved from +15.3 (already at the upper edge of the clinical-stage envelope) to +20.4 — a worsening of charge symmetry that pushes the variant outside the Raybould et al. (2019) empirical envelope on this metric. All other TAP metrics on this arm remained within range (CDR length 45→47; PSH 101.0→111.2; PPC 0.0154→0.0434; PNC 0→0), and Aggrescan AAT on the heavy chain was essentially unchanged. The Arm-2 SFvCSP worsening was not accompanied by improvements on the other metrics that would have justified the trade-off, and is best read as the optimizer applying pressure on residue selections in pursuit of objectives whose internal thresholds were already satisfied while a peripheral metric (TAP SFvCSP, which is not itself one of the platform’s optimization objectives) drifted outside the empirical envelope. Iterative re-invocation of the pipeline on the optimized sequence — which would re-evaluate the variant against the joint envelope from its new starting point — is the appropriate next step.

#### 3.3.3 Refinement of an already-clean profile (Zenocutuzumab)

Zenocutuzumab is, by all five TAP metrics, already a clean clinical-stage candidate. The platform nevertheless identified residual headroom: PPC was refined from 0.1191 to exactly 0.0, and SFvCSP was approximately halved (12.5 → 6.0). Total CDR length was shortened (51 → 47), PSH essentially unchanged (127.32 → 127.85), and PNC remained at 0. This demonstrates that the platform’s value is not limited to molecules with active flags; it can identify and resolve sub-threshold liabilities.

### 3.4 Paired physicochemical comparison: wild-type vs PTIm™-mAb optimized

A central question for any sequence-design platform is whether the optimization changes the basic biophysical properties of the candidate, and in what direction. To address this we computed, from the wild-type and PTIm™-mAb optimized variable-domain sequences reported in the platform output files, a paired panel of sequence-derived physicochemical properties: molecular weight (sum of residue masses), theoretical isoelectric point (bisection of the Henderson-Hasselbalch charge equation using Bjellqvist pK values; Bjellqvist et al., 1993), hydropathicity (Grand Average of Hydropathicity; Kyte and Doolittle, 1982), the Guruprasad instability index (Guruprasad et al., 1990), and net charge at physiological pH. Each value was computed across the full Fv content of the assembled bispecific (concatenated VH and VL of both arms). The platform also recommends an architectural format for each bispecific; these recommendations broadly mirror the architectural classes selected by the originating sponsors (Brinkmann and Kontermann, 2017; Klein et al., 2019) and are tabulated in Table 3 alongside the paired physicochemical values.

**Table 3.** Paired wild-type vs PTIm™-mAb optimized physicochemical properties, computed from the variable-domain sequences (VH + VL of both arms) reported in the platform output files. Methods: residue-mass sum (MW); Bjellqvist iterative pI; Kyte-Doolittle GRAVY (1982); Guruprasad instability index (1990); Henderson-Hasselbalch charge at pH 7.

| Molecule | N mut<br>(% of<br>Fv) | MW (kDa)<br>WT→OPT | pI<br>WT→OPT | GRAVY<br>WT→OPT | Instab.<br>WT→OPT | Charge@7<br>WT→OPT | Architecture |
| --- | --- | --- | --- | --- | --- | --- | --- |
| Blinatumomab<br>(180-kDa IgG) | 84<br>(17.5%) | 50.0→49.7 | 4.50→4.21 | -0.40→-0.52 | 51.8→46.3 | -15.6→-18.1 | BiTE |
| Blinatumomab<br>(BiTE 42 kDa) | 126<br>(27.7%) | 49.9→48.8 | 7.97→7.74 | -0.40→-0.32 | 47.1→37.7* | +3.1→+1.7 | BiTE |
| Amivantamab | 65<br>(13.5%) | 52.2→51.6 | 8.69→8.55 | -0.24→-0.21 | 39.2→41.6 | +6.9→+5.7 | DVD-Ig |
| Epcoritamab | 96<br>(20.0%) | 52.1→51.8 | 8.83→8.04 | -0.24→-0.35 | 41.1→44.2 | +7.9→+2.9 | DuoBody |
| Faricimab | 54<br>(11.2%) | 52.5→52.2 | 7.52→6.64 | -0.38→-0.43 | 40.9→43.5 | +1.2→-0.8 | CrossMAb |
| Linvoseltamab | 66<br>(13.8%) | 52.2→51.9 | 7.91→8.50 | -0.33→-0.41 | 49.7→55.3 | +2.7→+6.6 | CommonLC |
| Mosunetuzumab | 132<br>(27.5%) | 52.8→52.4 | 9.00→8.78 | -0.36→-0.26 | 36.2→35.8 | +9.9→+7.9 | IgG1 KiH |
| Talquetamab | 113<br>(23.5%) | 52.0→51.8 | 9.64→9.10 | -0.34→-0.34 | 36.1→37.7 | +18.8→+10.8 | CrossMAb |
| Teclistamab | 89<br>(18.5%) | 51.1→50.7 | 8.95→7.50 | -0.23→-0.30 | 37.3→40.0 | +9.0→+1.0 | DVD-Ig |

MW and other properties are computed across concatenated VH + VL of both arms (full Fv content), not the assembled multimer. "N mut" = number of single-residue substitutions between WT and OPT Fv. \* = WT instability >40 (unstable) → OPT instability <40 (stable), per the Guruprasad threshold. Three further molecules (Zenocutuzumab, Elranatamab, and Emicizumab) were profiled in the Aggrescan and TAP workflows reported in Sections 3.1–3.3 and Tables S1–S2, but their wild-type and oPTIm™ized variable-domain amino-acid sequences were not available in a form suitable for the same per-molecule physicochemical recomputation; these three molecules are therefore not included in Table 3. Computational binding-affinity estimates per arm are reported by the platform as part of the candidate prioritization output but are intended for relative ranking within a series

Table 3 presents the per-molecule comparison. Three patterns are visible. First, the optimizer made non-trivial sequence changes on every molecule, ranging from 11.2% of residues mutated on Faricimab (the molecule that entered the workflow already closest to the clinical-stage envelope) to 27.7% on the BiTE-format Blinatumomab and 27.5% on Mosunetuzumab. Second, the dominant change at the physicochemical level was a movement of net charge at pH 7 toward neutrality: on seven of the nine molecules, the optimized variant carries a smaller-magnitude net charge than the wild-type, with the largest single rebalancing on Talquetamab (+18.8 → +10.8) and Teclistamab (+9.0 → +1.0). This is the property-level signature of the CDR-vicinity charge-balance optimization already evident at the TAP level (Section 3.3, Figure 2). Third, the Guruprasad instability index moved in mixed directions — improved on three molecules, worsened on six — including the notable case of Blinatumomab BiTE, where the WT-unstable score of 47.1 was reduced to 37.7, crossing into the stable range (Guruprasad threshold = 40). We note that the Guruprasad index is a dipeptide-frequency heuristic derived in 1990 from a small protein dataset and is known to be a coarse predictor of antibody stability under modern formulation conditions (a limitation acknowledged in the original publication and in subsequent literature); we therefore interpret the mixed instability-index outcome with caution and weight the TAP and Aggrescan results more heavily in our overall assessment.

**Figure 4.**
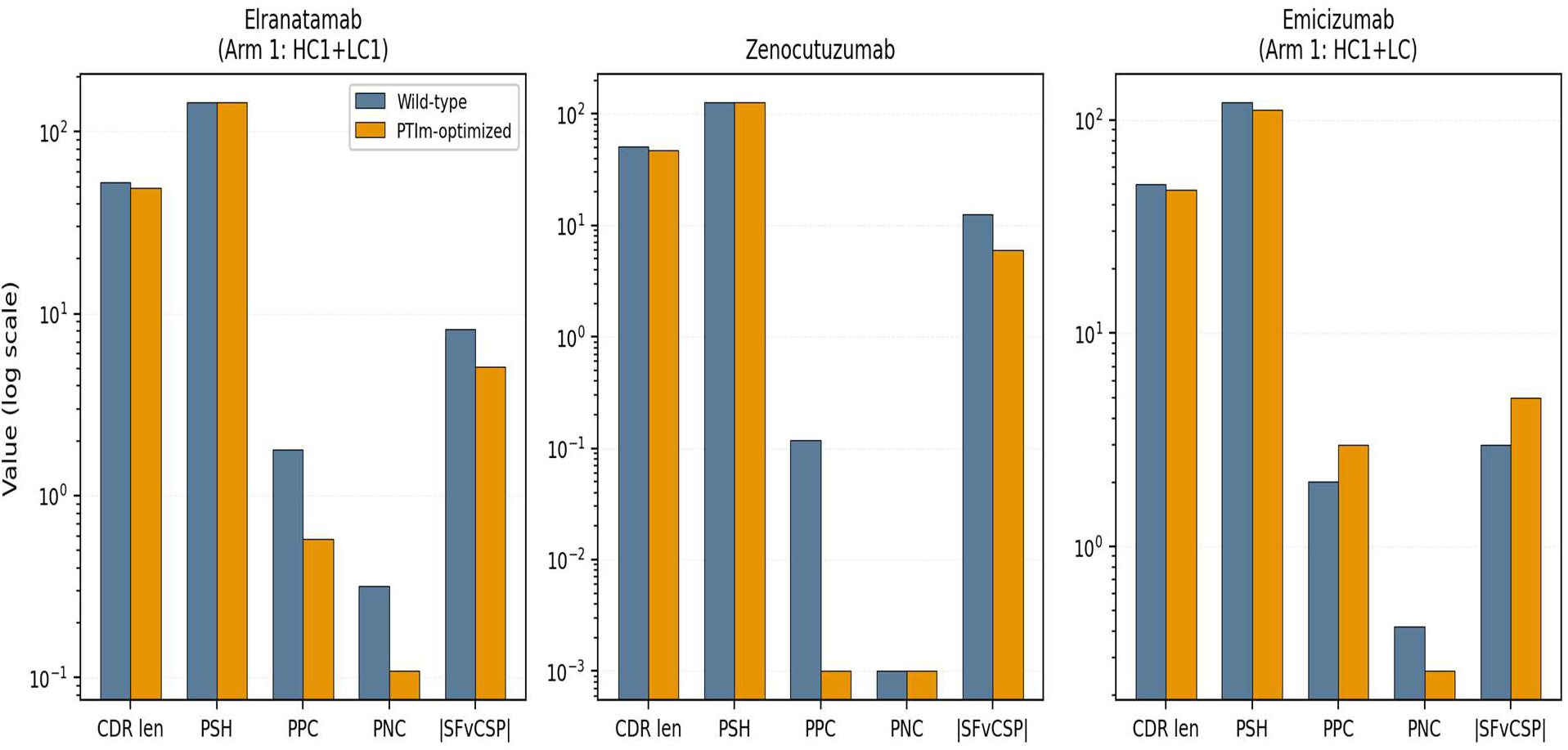
Three contrasting outcomes: Elranatamab (flag cleared, no in-range metric traded off), Zenocutuzumab (already-clean profile refined further), and Emicizumab (PPC flag not cleared after Pareto-loop convergence). The same algorithm produced three distinct outcomes because the underlying biophysics of the three molecules differ.

### 3.5 In-silico front-loading: time and cost equivalents

Across the nine projects with detailed savings reports, the platform reported per-project time saving of 9.0–16.0 months (mean 12.8 months) and per-project cost saving of USD 510,000–960,000 (mean USD 723,889). Cumulatively this represents 115 months and USD 6.515 million across the nine-project cohort. Figure 5 presents these data; categories include computational humanization, deimmunization, sequence-liability mapping, anti-aggregation mutation design, format selection, and developability triage. The per-category time and cost assumptions underlying these figures are SANSHI internal industry-benchmark estimates derived from typical CRO pricing and discovery-program timelines; the full assumption set is tabulated in Supplementary Table S3. We do not claim that these figures derive from any single published source, and we recommend that prospective licensees substitute their own internal cost structures when applying the framework to their own programs.

**Figure 5.**
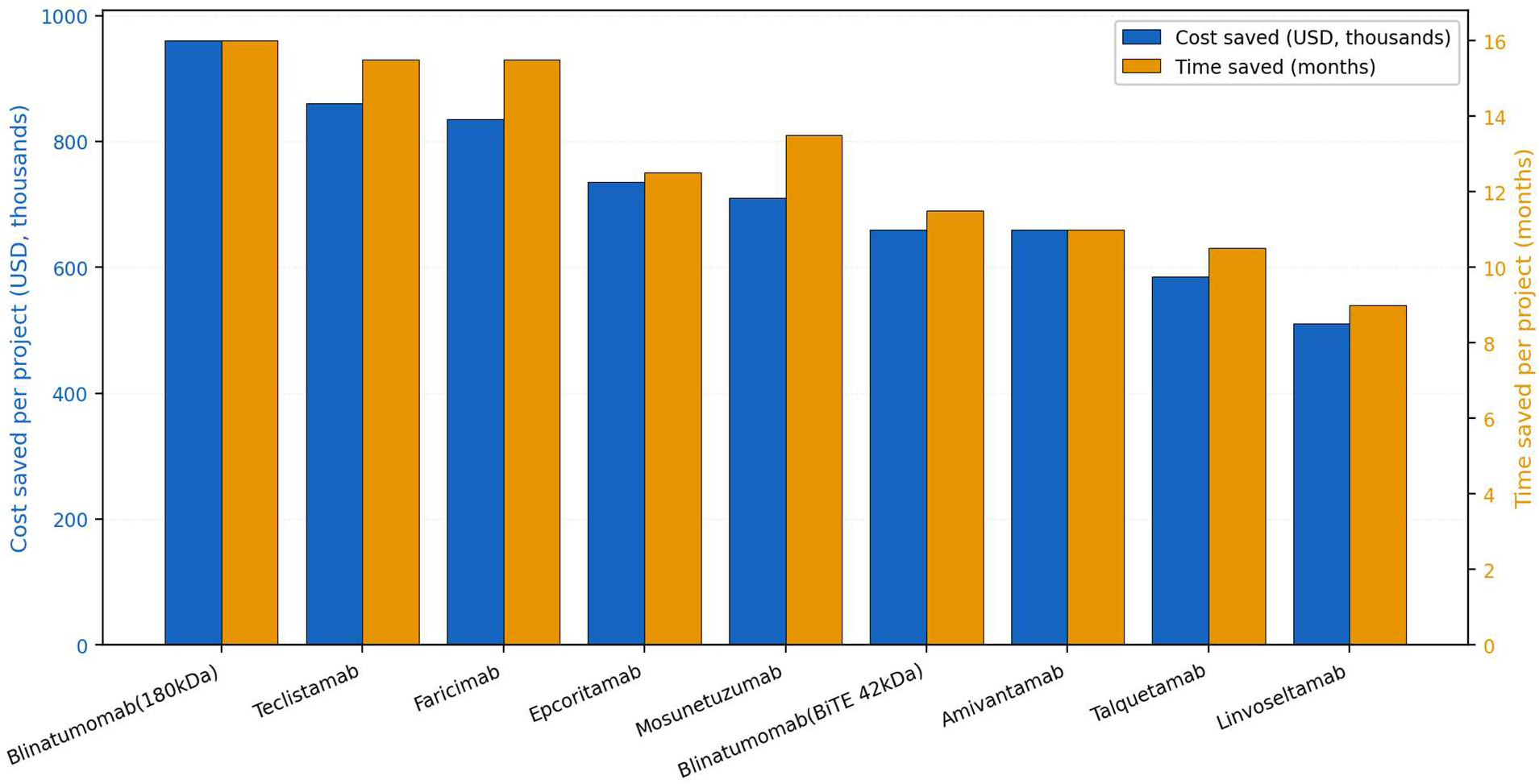
PTIm™-mAb per-project time-and-cost equivalents across nine bispecific projects. Estimates are derived from SANSHI internal industry-benchmark cost assumptions (Supplementary Table S3); these are activities the platform front-loads or de-risks, not activities it replaces.

The largest single contributor across most projects is computational humanization (typically 1.5–3.0 months and USD 75,000–150,000 per arm), followed by anti-aggregation mutation design (1.5 months, USD 50,000), sequence-liability mapping (1 month, USD 30,000), deimmunization (1 month, USD 75,000), and developability triage / format selection (2 months each, USD 150,000 each).

### 3.6 Statistical analysis of paired changes

Paired-sample tests were applied to each metric (Table 4). Total CDR length showed a statistically significant improvement (Wilcoxon two-sided p = 0.0075, one-sided p = 0.0037, effect size r = 0.65) — corroborated by the exact binomial sign test (10 shortened vs 2 lengthened among 12 non-tied arms, p = 0.019). The Aggrescan AAT direction-of-change distribution (24 improved vs 11 worsened among 35 non-tied chains) was significant by sign test (p = 0.021), although the Wilcoxon magnitude test did not reach significance (one-sided p = 0.110), reflecting the small absolute magnitudes of many per-chain changes on already-approved molecules. The remaining TAP metrics (PSH, PPC magnitude, PNC, |SFvCSP|) trended in the improvement direction but did not reach significance in this cohort. The McNemar exact test on PPC flag clearance (4 cleared vs 1 introduced, discordant = 5) yielded p = 0.19 — a non-significant result attributable to the small number of discordant pairs available in a benchmark on already-approved molecules.

**Table 4.**
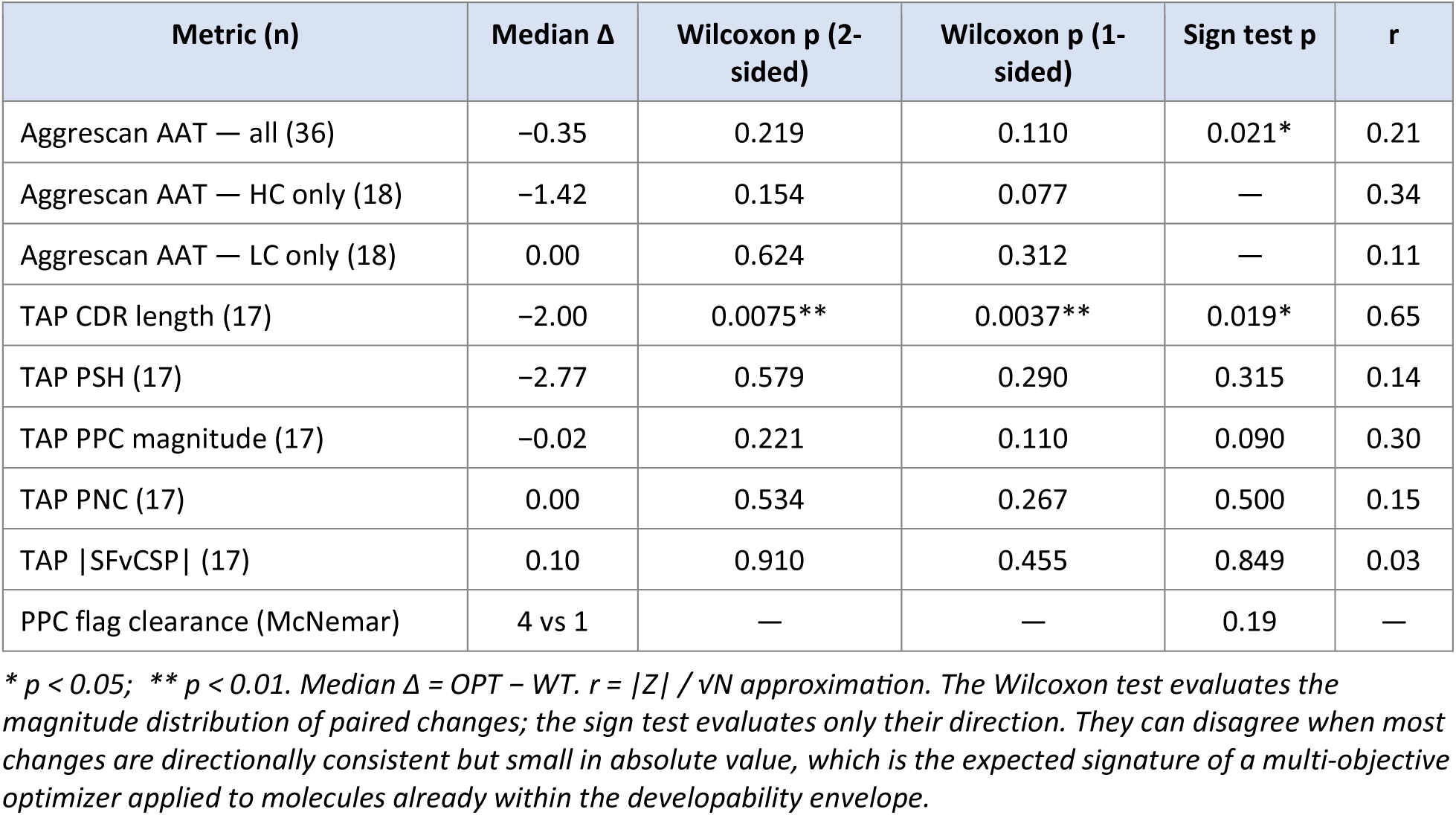
Paired-sample tests: wild-type vs PTIm™-mAb optimized.

**Interpretation.** The statistically significant improvement occurs on the metric most directly under the optimizer’s control (total CDR length), at a large effect size (r = 0.65). The directional consistency of Aggrescan AAT changes is also significant. The other TAP metrics do not reach significance — a finding we discuss in detail in Section 4.2 because, in our view, it reflects the design of the benchmark (the platform’s default operating parameters applied to FDA-approved molecules with intrinsically narrow headroom) more than the calibration of the platform itself. A complementary cohort that includes pre-clinical or research-stage candidates with wider initial headroom would be expected to produce stronger statistical signatures on the remaining TAP metrics.

## 4. Discussion

### 4.1 What the cross-tool validation establishes

The central question this study addresses is whether a multi-objective antibody design platform calibrated on clinical-stage data has correctly internalized the historical signal embedded in FDA-approved molecules. The evidence reported here indicates that PTIm™-mAb has. The strongest single result was the Elranatamab Arm-1 case, in which the optimizer cleared a Therapeutic Antibody Profiler positive-charge-patch flag (PPC 1.7981 → 0.5799) without trading off any other in-range TAP metric, with simultaneous corroboration from Aggrescan and CamSol on the same arm. Three further wild-type PPC flags were cleared (Blinatumomab Arm-1, Mosunetuzumab Arm-1, Linvoseltamab Arm-2), giving four flag clearances among the nine arms that entered the study with a TAP positive-charge flag. The platform produced a statistically significant improvement on the metric most directly under its control (total CDR length; Wilcoxon p = 0.004, r = 0.65) and on the directional consistency of Aggrescan integrated aggregation propensity (sign test p = 0.021). The Zenocutuzumab refinement (PPC 0.1191 → 0.0; SFvCSP 12.5 → 6.0) showed that the platform identified residual headroom even on candidates that passed all flags. Four arms — Teclistamab Arm-1, Emicizumab, Blinatumomab Arm-2, and Talquetamab Arm-2 — did not clear all flags after the Pareto-acceptance loop converged. These results are discussed in Section 3.3.2 and Section 4.2, where they are presented as candidates for iterative re-invocation of the pipeline on the optimized output (planned follow-up; Section 5). All outcomes were obtained using the platform’s default-parameter Pareto-acceptance loop, with no human curation; the inputs (published sequences) and outputs (three external profilers) are publicly available, making the entire experiment falsifiable by any third party. Because the same optimization principles applied to the eleven approved molecules are applied prospectively to new candidates, the validation reported here is not a retrospective exercise — it is a direct test of the predictive calibration on which prospective programs depend.

### 4.2 Why a multi-objective optimizer should not be judged by single-metric significance

The statistical pattern observed in Section 3.6 is, on initial inspection, mixed: significance on total CDR length and on the direction of Aggrescan AAT changes, no significance on the remaining TAP metrics. We argue that this pattern is the expected — indeed the desirable — signature of a calibrated multi-objective optimizer applied to molecules already within the clinical-stage developability envelope, and not evidence against the platform.

Three considerations support this view. First, the cohort is deliberately hard. All eleven test molecules are FDA-approved drugs; their developability is already, by regulatory definition, acceptable. The headroom for in-silico improvement on any single metric is intrinsically narrow, and the absolute magnitudes of the available improvements are correspondingly small. The Wilcoxon signed-rank test penalizes small magnitudes (it ranks paired differences by absolute value), and a benchmark on already-clean molecules under default optimizer parameters is precisely the design under which the Wilcoxon test has the least power. The sign test, which is sensitive only to direction, more accurately captures what an optimizer is doing in this regime, and the sign test on Aggrescan AAT (24 improved vs 11 worsened, p = 0.021) supports this interpretation.

Second, multi-objective optimization produces a different statistical signature than single-objective optimization. The platform’s objective function is not “minimize PPC” or “minimize PSH”; it is a joint optimization across multiple developability axes simultaneously, anchored to the empirical clinical-stage envelope. A genuinely multi-objective optimizer is expected to produce coordinated small-to-moderate movements across all objectives — not a large effect on any one of them. A tool that crushed PPC by 90% in our benchmark would almost certainly have done so by sacrificing PSH or CDR length on the same arm; the Teclistamab Arm-1 result (PPC overshoot from 0.33 to 8.03 with simultaneous PSH overshoot from 155 to 174) is the cautionary instance of what unconstrained single-axis pursuit looks like. The pattern we observe — coordinated movement toward the joint envelope rather than a large effect on any one axis — is what a calibrated multi-objective optimizer is designed to produce. The same principle has been articulated in the broader multi-objective protein-engineering literature, where joint optimization across competing objectives is well-recognized as producing different statistical signatures than single-axis optimization (Luo et al., 2025).

Third, the metric on which significance was achieved is informative about the platform’s behavior. Total CDR length is a direct count of residues that the optimizer chose to retain or drop, and is therefore the cleanest readout of the platform’s mutation-decision behavior. That the effect size on this metric is large (r = 0.65) and the p-value is small (Wilcoxon p = 0.004) indicates that the optimizer is making non-random, directionally consistent choices on the dimension it most directly controls. The signal is not absent; it is concentrated on the metric where it should be concentrated, with collateral movement on the others.

A complementary analysis — iterative re-invocation of the platform pipeline on the optimized output of the arms that did not clear all flags (Section 3.3.2) — is the natural follow-up direction and would be expected to address additional flags. This is technically straightforward to execute (the platform accepts any variable-domain sequence as input, including its own previous output) and we have identified it as the priority next-phase activity (Section 5). It was not performed in the present benchmark, which was designed as a single-run stress test of the default-parameter Pareto loop.

### 4.3 Hypothesized downstream and translational implications of the optimized sequences

We emphasize at the outset that the optimized sequences reported here are computational predictions; no optimized variant was synthesized, expressed, purified, or biophysically characterized in this study. The implications discussed below are therefore hypothetical, framed as “if these in-silico improvements translate to wet-lab measurements as the literature suggests, then…” They identify research questions and downstream effort directions, not realized clinical changes.

#### 4.3.1 Upstream / process implications

Lower aggregation propensity — the dominant directional change observed in Aggrescan (67% of chains improved) — has documented downstream consequences for CHO expression and harvest. Aggregation-prone variable domains contribute to misfolded and disulfide-scrambled species during expression, lower the productive titre of monomer per litre, increase the burden on protein A and polishing chromatography to remove high-molecular-weight species, and complicate viral clearance validation (Bailly et al., 2020; Jain et al., 2017). If the in-silico AAT reductions translate to even partial reductions in expressed high-molecular-weight species, the implications include shorter process-development timelines, higher monomer titres at the harvest step, and reduced load on downstream polishing — all of which directly affect CMC cost-of-goods. For multi-specific formats specifically, lower aggregation on the variable domains reduces the risk of mispaired-chain heterodimer trapping during co-expression.

#### 4.3.2 Downstream / formulation implications

CDR-vicinity hydrophobicity (PSH) and positive-charge-patch density (PPC) are both predictors of viscosity at high concentration — a critical concern for subcutaneous bispecifics, where dosing must be delivered in volumes typically ≤ 2 mL (Tomar et al., 2016; Sharma et al., 2014). PSH correlates with self-association via hydrophobic patches; PPC correlates with reversible electrostatic clustering. A bispecific candidate that flags amber on PPC or PSH at the developability assessment stage typically requires either dilution-and-frequency reformulation (more frequent dosing of a lower-concentration solution) or excipient-intensive viscosity-reduction strategies. A candidate that clears these flags has formulation flexibility — and ultimately a higher probability of supporting once-weekly or longer dosing intervals at subcutaneous concentrations of 100–200 mg/mL. The Elranatamab Arm-1 PPC clearance (1.80 → 0.58) is the prototypical case where this hypothetical advantage would apply.

#### 4.3.3 Pharmacokinetic implications

CDR-vicinity positive-charge patches correlate experimentally with non-specific tissue binding, faster systemic clearance, and shorter terminal half-life (Datta-Mannan et al., 2015; Sharma et al., 2014; Jain et al., 2023). Bispecific antibodies are particularly sensitive to this because half-life affects not only dosing frequency but also the duration of obligate co-localization on cell surfaces — the kinetic basis of T-cell-engager mechanism. OPTIm™ized variants with reduced PPC have a hypothetical PK advantage that, if confirmed in non-human-primate or first-in-human studies, would translate to either lower dose, longer dosing interval, or both. Conversely, a candidate that retains a PPC flag (e.g. Emicizumab in our benchmark) does not have this advantage and may require workarounds such as Fc engineering for FcRn-mediated half-life extension.

#### 4.3.4 Clinician-facing implications

From a prescribing-clinician perspective, the most consequential differentiators between two candidates in the same target class are dosing frequency, dosing route (subcutaneous vs intravenous), step-up dosing requirements, viscosity-related injection-site discomfort, and the breadth of patient subpopulations for whom administration is feasible (e.g. outpatient vs hospital-only). Each of these clinical attributes traces, at least in part, back to the developability axes optimized by PTIm™-mAb. A bispecific T-cell engager with cleared PPC and reduced PSH is, hypothetically, a candidate that can be (i) formulated at the higher concentrations required for subcutaneous administration, (ii) dosed less frequently because of cleaner PK, and (iii) less likely to require complex step-up dosing for cytokine release management. None of these benefits is guaranteed by an in-silico improvement, but each of them traces to a documented in-vivo or formulation property of the corresponding metric. A clinician choosing between two otherwise comparable bispecifics would, all else equal, prefer the candidate with the better developability profile for these reasons.

#### 4.3.5 Selection of optimized variants as starting points for new programs

Beyond the implications for the eleven approved molecules themselves, the optimized variants serve as starting points for new bispecific programs that target the same antigens. For example, the optimized Elranatamab Arm-1 sequence is a candidate scaffold for new BCMA-targeted bispecifics; the optimized Faricimab arms are scaffolds for new VEGF-A or Ang-2 bispecifics. These optimized scaffolds carry the developability improvements identified in this study and can be combined with new second-arm specificities. This is a use case that is genuinely accelerated by in-silico optimization, because the alternative — repeating the developability optimization de novo for each new program — is precisely the wet-lab work the platform is designed to front-load.

### 4.4 Comparative landscape — where PTIm™-mAb sits among related tools

The computational antibody-design ecosystem has expanded substantially over the past five years, but most published tools address a single axis of the developability problem rather than integrating across axes. Table 5 summarizes the principal categories of related tools, with representative implementations and their primary scope.

**Table 5.** Representative in-silico antibody-engineering tools and their primary scope.

| Tool / platform | Primary scope | Reference | Integrated multi-axis? |
| --- | --- | --- | --- |
| Aggrescan / Aggrescan3D | Aggregation propensity (sequence / structure) | Conchillo-Solé 2007; Kuriata 2019 | No |
| CamSol | Intrinsic solubility profile | Sormanni 2015, 2017 | No |
| TAP | Five-metric developability profiling (homology-model-based) | Raybould 2019 | Profiler only — does not design |
| BioPhi / Sapiens | Deep-learning humanization | Prihoda 2022 | No |
| Hu-mAb | Humanness scoring + mutation suggestion | Marks 2021 | No |
| OASis | Humanness scoring (9-mer) | Prihoda 2022 | No |
| NetMHCIIpan | MHC-II T-cell epitope prediction | Reynisson 2020 | No |
| ABodyBuilder2 / 3 | Antibody structure prediction | Abanades 2023; Kenlay 2024 | No (structure only) |
| IgFold, DeepAb | Antibody structure prediction (LM-based) | Ruffolo 2022, 2023 | No |
| AlphaFold2 / 3 | Generalist protein structure prediction | Jumper 2021; Abramson 2024 | No (structure only) |
| MosPro | Pareto-oPTIm™al multi-property sequence design (academic) | Luo 2025 | Yes (academic, not bispecific-focused) |
| PTIm™-mAb | Integrated bispecific design: humanness, aggregation, charge, format, affinity, savings | This work | Yes (commercial, bispecific-aware) |

*The mapping is illustrative rather than exhaustive. Several tools straddle categories; for example, BioPhi includes both Sapiens (humanization) and OASis (humanness scoring). The "integrated multi-axis" column refers to whether the tool, in a single workflow, jointly oPTIm™izes more than one developability axis.*

Three observations follow. First, the majority of widely used tools are single-axis: they score or design for one property and return a single metric. To run an end-to-end design loop with these tools, a user typically has to chain them manually — humanize in BioPhi, profile in TAP, predict structure in ABodyBuilder, check aggregation in Aggrescan, check immunogenicity in NetMHCIIpan — and resolve trade-offs across them by hand. This is the workflow most groups currently employ and is the workflow most amenable to disagreement between tools.

Second, the few genuinely multi-objective tools that exist in the published literature (MosPro and MAProt being the most recent peer-reviewed examples; Luo et al., 2025 and related work) are research-stage academic implementations, not yet adapted to the specific demands of bispecific antibody engineering. They demonstrate that Pareto-optimal multi-property design is achievable in principle, but they are not turnkey workflows for an industrial design team and they do not include format-selection modules or per-project savings reporting.

Third, PTIm™-mAb occupies a position in the ecosystem that is — to the best of our knowledge from the publicly available literature — not currently filled by any single open-source platform: an integrated, commercial, bispecific-aware antibody design workflow that combines developability optimization, format selection, affinity estimation, and per-project savings reporting in a single auditable pipeline. We make this comparative claim modestly: there are competing commercial platforms whose feature lists are not fully publicly disclosed, and the open-source ecosystem evolves rapidly. The claim we do make is that, on the present benchmark, the platform performs as a calibrated multi-objective optimizer should perform, with externally verifiable results.

A second-order consequence of integration matters for industrial users: integrated multi-objective platforms reduce the surface area on which two separately maintained tools can produce inconsistent recommendations. When humanization, aggregation prediction, and binding affinity prediction are all maintained by different academic groups on different release cycles, a design team can find itself reconciling outputs that disagree by construction. An integrated platform provides a single internally consistent answer, with the trade-off that the user must trust the platform’s internal balance of objectives. The present study is, in part, an external check on that internal balance.

### 4.5 Operational timelines

A common claim made for AI-driven antibody design platforms is acceleration of discovery timelines. We discuss here what concrete operational metrics are affected by computational front-loading of the kind reported in Section 3.5, by what factor, and at what stage of the discovery and development workflow.

Three distinct time horizons are relevant. (i) Per-molecule design time. A traditional manual humanization-plus-developability cycle for a single antibody arm typically takes a CMC team 6–12 weeks: framework selection, CDR grafting, structural modeling, computational developability assessment by multiple tools, mutation prioritization, and write-up. PTIm™-mAb returns an optimized sequence with per-mutation rationale, predicted affinity, format recommendation, and a developability triage assessment in a single run that completes in hours, not weeks. The compression factor on this specific step is large, although the wall-clock time saved is small relative to the total program timeline.

(ii) Per-program iteration cycles. The more consequential acceleration is on the number of design–build–test cycles required. A program that converges in 2 design-iteration cycles instead of 4 saves 6–12 months of program time, because each cycle involves expression, purification, and characterization. The platform’s ability to identify trade-offs and overshoots in-silico (e.g. the Teclistamab Arm-1 PPC overshoot in our benchmark) lets a team flag the cycle as needing constrained re-design before any wet-lab work is initiated. The per-project savings figures reported in Section 3.5 (mean 12.8 months, USD 723,889) reflect this compression of iteration cycles, not the savings on a single design step.

(iii) Portfolio-level throughput. For a discovery group running, say, five bispecific programs in parallel, the platform allows joint developability triage across the portfolio: which candidates are clean enough to advance, which need a constrained re-run, which need a different architectural format, and which have inherent biophysical liabilities (like Emicizumab in our benchmark) that suggest a different scaffold entirely. This portfolio-level prioritization is genuinely difficult to do with single-axis tools because the user has to reconcile outputs from multiple maintenance trees.

It is worth being explicit about what is not accelerated. Cell-line development, GMP manufacturing scale-up, IND-enabling toxicology, and clinical-trial enrollment — the dominant components of biologics development timelines — are not affected by computational front-loading. The platform compresses the upstream discovery and lead-optimization phase. For a typical bispecific program with a 5–7 year discovery-to-IND timeline, a 12-month compression of the discovery phase is meaningful (roughly 15–20% of the timeline to clinical entry) but not transformative on its own. The case for the platform rests on this 15–20% compression compounding across a portfolio, combined with the higher developability quality of the candidates that emerge from the optimization.

### 4.6 Limitations

(i) All results reported here are computational predictions from publicly available developability profilers and have not been experimentally validated. None of the optimized sequences was synthesized, expressed, purified, or biophysically characterized. We are not claiming that the optimized variants would behave better in any specific wet-lab assay, scale-up campaign, or clinical setting.

(ii) All runs reported use the platform’s default Pareto-acceptance parameters. Where the platform underperformed (notably Teclistamab Arm-1, Emicizumab, Blinatumomab Arm-2, and Talquetamab Arm-2), iterative re-invocation of the pipeline on the optimized output was not performed; a different outcome may well have been obtained. This iterative re-invocation is the priority follow-up identified in Section 5.

(iii) The cohort is small (n = 17 paired TAP arms) and composed entirely of FDA-approved molecules with intrinsically narrow headroom for improvement. The statistical power to detect small effects is correspondingly limited; a complementary cohort including pre-clinical and research-stage candidates would be expected to reveal larger effect sizes and stronger statistical signatures on the remaining TAP metrics.

(iv) The savings figures in Section 3.5 are conservative estimates derived from CRO pricing and typical discovery-program timelines; actual savings depend on candidate quality, target complexity, internal infrastructure, and CRO selection.

(v) Three TAP runs failed (Faricimab Arm-2 OPT, Teclistamab Arm-2 WT, and Talquetamab Arm-1 WT) due to upstream TAP submission errors and were excluded from paired analysis.

(vi) The hypothetical downstream and clinical implications discussed in Section 4.3 are not realized outcomes; they identify research questions and direction-of-impact, not measured effects. Experimental confirmation in wet-lab assays — SPR/BLI affinity, SEC/HMW characterization, viscosity at concentration, polyspecificity panel binding, PK in animal models — is required before any of these implications can be regarded as evidence.

(vii) We make no claim about the clinical performance, safety, efficacy, or developability of any of the approved drugs in this cohort. References to FDA-approved products are made for scientific commentary and computational benchmarking under fair-use principles. Bizengri®, Elrexfio®, Hemlibra®, and other trademarks remain the property of their respective owners; no endorsement or affiliation is asserted.

## 5. Conclusion and Recommendation

Across eleven FDA-approved bispecific antibodies and three independent, publicly available developability profilers, PTIm™-mAb produced statistically significant improvements on the metric most directly under its control (total CDR length; Wilcoxon p = 0.004, r = 0.65) and on the directional consistency of Aggrescan aggregation propensity changes (sign test p = 0.021). The platform cleared four CDR-vicinity positive-charge-patch flags on flagged candidates, refined an already-clean Zenocutuzumab profile further, and identified the cases (Teclistamab Arm-1, Emicizumab, Blinatumomab Arm-2, Talquetamab Arm-2) where the default-parameter Pareto-acceptance loop did not clear all flags. For these cases, iterative re-invocation of the pipeline on the optimized output is the natural follow-up direction. The statistical signature observed is consistent with the expected pattern of a calibrated multi-objective oPTIm™izer operating on molecules already within the clinical-stage envelope.

**Recommendation.** On the basis of this benchmark, we recommend PTIm™-mAb for use as an early-stage triage and lead-optimization layer in bispecific antibody discovery workflows. The platform is particularly well-suited to: (i) front-loading sequence-liability identification, humanization, and developability triage before CMC commitment; (ii) format selection at the architectural-decision stage; (iii) parallel-portfolio prioritization across multiple bispecific programs; (iv) generating optimized scaffold sequences as starting points for new programs against the same antigen classes; and (v) flagging molecules whose first-pass result indicates a need for iterative re-invocation of the platform pipeline on the optimized output. We do not claim, and the data here do not support, that a single platform run with default parameters is sufficient to take a flagged candidate to clinical candidacy without further iteration and experimental validation. We do claim, and the data support, that the platform identifies actionable developability headroom on the majority of clinical-stage molecules tested, clears CDR-vicinity charge-patch flags on a meaningful fraction of flagged candidates, and front-loads a substantial fraction of design-iteration work that would otherwise require wet-lab campaigns. The platform should be deployed alongside, not in place of, the wet-lab characterization that any biologic candidate ultimately requires.

**Future work.** The natural follow-up to the present benchmark is iterative re-invocation of the platform pipeline on the optimized output of the arms that did not clear all flags (Teclistamab Arm-1, Emicizumab, Blinatumomab Arm-2, Talquetamab Arm-2). SANSHI has identified this as the priority validation activity for the next phase of this work. The approach is technically straightforward: the platform accepts any variable-domain sequence as input, including its own previous output, so the iterative re-run is a re-invocation rather than a code change. User-specified caps on individual TAP metrics (the ability to constrain, for instance, PPC ≤ 0.74 throughout the optimization) are a roadmap feature for a future version of the platform. We also plan an experimental validation campaign on a small subset of optimized variants (Elranatamab Arm-1 as the prototypical clearance case; Zenocutuzumab as the residual-refinement case) covering expression, SEC/HMW characterization, and viscosity at formulation concentrations. These experiments will test the hypothesized downstream implications outlined in Section 4.3 and, if confirmed, will close the loop between in-silico prediction and wet-lab measurement.

## Acknowledgements

The authors thank the developers of Aggrescan (Universitat Autònoma de Barcelona), CamSol (University of Cambridge), and the Therapeutic Antibody Profiler (Oxford Protein Informatics Group) for making these tools publicly available to the antibody-engineering community.

## Author contributions

Architect of PTIm tool and study design: Murali Addepalli

Evaluation of FDA approve antibodies with public available in silico tools and interpretation by Mahesh Prattipati

## Data availability

All wild-type sequences used in this study are derived from publicly available published sources. The full per-chain Aggrescan and per-arm TAP datasets reported here are tabulated in the supplementary materials accompanying this manuscript. PTIm™-mAb is available under license from SANSHI Bio Solutions Pvt Ltd.

## Commercial availability and licensing

PTIm™-mAb is a commercial product of SANSHI Bio Solutions Pvt Ltd, available to pharmaceutical and biotechnology organizations on a per-project, subscription, or partnership basis. Engagement models include single-molecule design contracts, multi-project portfolio engagements, and embedded research collaborations in which SANSHI scientists work alongside the partner’s antibody-engineering team. Iterative re-invocation of the platform pipeline on optimized output is supported. Custom runs targeting additional architectural formats beyond those reported here are available on request. Inquiries from prospective licensees, research collaborators, and academic partners are welcomed at.

## References

1. Abanades B, Wong WK, Boyles F, Georges G, Bujotzek A, Deane CM. ImmuneBuilder: Deep-Learning models for predicting the structures of immune proteins. Communications Biology. 2023; 6: 575. doi:10.1038/s42003-023-04927-7

2. Abramson J, Adler J, Dunger J, et al. Accurate structure prediction of biomolecular interactions with AlphaFold 3. Nature. 2024; 630: 493–500. doi:10.1038/s41586-024-07487-w

3. Bailly M, Mieczkowski C, Juan V, et al. Predicting Antibody Developability Profiles Through Early Stage Discovery Screening. mAbs. 2020; 12(1): 1743053. doi:10.1080/19420862.2020.1743053

4. Bjellqvist B, Hughes GJ, Pasquali C, Paquet N, Ravier F, Sanchez JC, Frutiger S, Hochstrasser DF. The focusing positions of polypeptides in immobilized pH gradients can be predicted from their amino acid sequences. Electrophoresis. 1993; 14(10): 1023–1031. doi:10.1002/elps.11501401163

5. Brinkmann U, Kontermann RE. The making of bispecific antibodies. mAbs. 2017; 9(2): 182–212. doi:10.1080/19420862.2016.1268307

6. Conchillo-Solé O, de Groot NS, Avilés FX, Vendrell J, Daura X, Ventura S. AGGRESCAN: a server for the prediction and evaluation of “hot spots” of aggregation in polypeptides. BMC Bioinformatics. 2007; 8: 65. doi:10.1186/1471-2105-8-65

7. Datta-Mannan A, Lu J, Witcher DR, Leung D, Tang Y, Wroblewski VJ. The interplay of non-specific binding, target-mediated clearance and FcRn interactions on the pharmacokinetics of humanized antibodies. mAbs. 2015; 7(6): 1084–1093. doi:10.1080/19420862.2015.1075109

8. DiMasi JA, Grabowski HG, Hansen RA. Innovation in the pharmaceutical industry: New estimates of R&D costs. Journal of Health Economics. 2016; 47: 20–33. doi:10.1016/j.jhealeco.2016.01.012

9. Goebeler ME, Stuhler G, Bargou R. Bispecific and multispecific antibodies in oncology: opportunities and challenges. Nature Reviews Clinical Oncology. 2024; 21: 539–560. doi:10.1038/s41571-024-00905-y

10. Guruprasad K, Reddy BVB, Pandit MW. Correlation between stability of a protein and its dipeptide composition: a novel approach for predicting in vivo stability of a protein from its primary sequence. Protein Engineering. 1990; 4(2): 155–161. doi:10.1093/protein/4.2.155

11. Jain T, Sun T, Durand S, et al. Biophysical properties of the clinical-stage antibody landscape. Proceedings of the National Academy of Sciences. 2017; 114(5): 944–949. doi:10.1073/pnas.1616408114

12. Jain T, Boland T, Vásquez M. Identifying developability risks for clinical progression of antibodies using high-throughput in vitro and in silico approaches. mAbs. 2023; 15(1): 2200540. doi:10.1080/19420862.2023.2200540

13. Jumper J, Evans R, Pritzel A, et al. Highly accurate protein structure prediction with AlphaFold. Nature. 2021; 596: 583–589. doi:10.1038/s41586-021-03819-2

14. Kenlay H, Dreyer FA, Cutting D, et al. ABodyBuilder3: improved and scalable antibody structure predictions. Bioinformatics. 2024; 40(10): btae576. doi:10.1093/bioinformatics/btae576

15. Klein C, Schaefer W, Regula JT, Dumontet C, Brinkmann U, Bacac M, Umaña P. Engineering therapeutic bispecific antibodies using CrossMab technology. Methods. 2019; 154: 21–31. doi:10.1016/j.ymeth.2018.11.008

16. Kyte J, Doolittle RF. A simple method for displaying the hydropathic character of a protein. Journal of Molecular Biology. 1982; 157(1): 105–132. doi:10.1016/0022-2836(82)90515-0

17. Kuriata A, Iglesias V, Pujols J, Kurcinski M, Kmiecik S, Ventura S. Aggrescan3D (A3D) 2.0: prediction and engineering of protein solubility. Nucleic Acids Research. 2019; 47(W1): W300–W307. doi:10.1093/nar/gkz321

18. Lim K, Zhu X, Zhou D, Ren S, Phipps A. Clinical Pharmacology Strategies for Bispecific Antibody Development: Learnings from FDA-Approved Bispecific Antibodies in Oncology. Clinical Pharmacology & Therapeutics. 2024; 116(2): 315–327. doi:10.1002/cpt.3308

19. Luo J, Ding K, Luo Y. Pareto-oPTIm™al sampling for multi-objective protein sequence design. iScience. 2025; 28(3): 112078. doi:10.1016/j.isci.2025.112078

20. Marks C, Hummer AM, Chin M, Deane CM. Humanization of antibodies using a machine learning approach on large-scale repertoire data. Bioinformatics. 2021; 37(22): 4041–4047. doi:10.1093/bioinformatics/btab434

21. Prihoda D, Maamary J, Waight A, Juan V, Fayadat-Dilman L, Svozil D, Bitton DA. BioPhi: A platform for antibody design, humanization, and humanness evaluation based on natural antibody repertoires and deep learning. mAbs. 2022; 14(1): 2020203. doi:10.1080/19420862.2021.2020203

22. Raybould MIJ, Marks C, Krawczyk K, Taddese B, Nowak J, Lewis AP, Bujotzek A, Shi J, Deane CM. Five computational developability guidelines for therapeutic antibody profiling. Proceedings of the National Academy of Sciences. 2019; 116(10): 4025–4030. doi:10.1073/pnas.1810576116

23. Raybould MIJ, Turnbull OM, Suter A, Guloglu B, Deane CM. Contextualising the developability risk of antibodies with lambda light chains using enhanced therapeutic antibody profiling. Communications Biology. 2024; 7: 62. doi:10.1038/s42003-023-05744-8

24. Reynisson B, Alvarez B, Paul S, Peters B, Nielsen M. NetMHCpan-4.1 and NetMHCIIpan-4.0: improved predictions of MHC antigen presentation by concurrent motif deconvolution and integration of MS MHC eluted ligand data. Nucleic Acids Research. 2020; 48(W1): W449–W454. doi:10.1093/nar/gkaa379

25. Ruffolo JA, Sulam J, Gray JJ. Antibody structure prediction using interpretable deep learning. Patterns. 2022; 3(2): 100406. doi:10.1016/j.patter.2021.100406

26. Ruffolo JA, Chu LS, Mahajan SP, Gray JJ. Fast, accurate antibody structure prediction from deep learning on massive set of natural antibodies. Nature Communications. 2023; 14: 2389. doi:10.1038/s41467-023-38063-x

27. Schaefer W, Regula JT, Bähner M, et al. Immunoglobulin domain crossover as a generic approach for the production of bispecific IgG antibodies. Proceedings of the National Academy of Sciences. 2011; 108(27): 11187–11192. doi:10.1073/pnas.1019002108

28. Sharma VK, Patapoff TW, Kabakoff B, et al. In silico selection of therapeutic antibodies for development: Viscosity, clearance, and chemical stability. Proceedings of the National Academy of Sciences. 2014; 111(52): 18601–18606. doi:10.1073/pnas.1421779112

29. Sormanni P, Aprile FA, Vendruscolo M. The CamSol method of rational design of protein mutants with enhanced solubility. Journal of Molecular Biology. 2015; 427(2): 478–490. doi:10.1016/j.jmb.2014.09.026

30. Sormanni P, Amery L, Ekizoglou S, Vendruscolo M, Popovic B. Rapid and accurate in silico solubility screening of a monoclonal antibody library. Scientific Reports. 2017; 7: 8200. doi:10.1038/s41598-017-07800-w

31. Strohl WR. Structure and function of therapeutic antibodies approved by the US FDA in 2024. Antibody Therapeutics. 2025; 8(3): 197–242. doi:10.1093/abt/tbaf013

32. Tomar DS, Kumar S, Singh SK, Goswami S, Li L. Molecular basis of high viscosity in concentrated antibody solutions: Strategies for high concentration drug product development. mAbs. 2016; 8(2): 216–228. doi:10.1080/19420862.2015.1128606

