## Supplementary material for "Benchmarking AI/ML-Driven PTIm-mAb Across Eleven FDA-Approved Bispecific Antibodies: A Cross-Tool Validation Study": Supplementary Data.docx

### Table S1. Full Aggrescan dataset — wild-type vs PTIm™-mAb optimized, all chains

| **Chain** | **WT nHS** | **WT NnHS** | **WT AAT** | **OPT nHS** | **OPT NnHS** | **OPT AAT** |
| --- | --- | --- | --- | --- | --- | --- |
| Blinatumomab_Arm1_HC | 7 | 5.833 | 13.895 | 6 | 5.000 | 12.411 |
| Blinatumomab_Arm1_LC | 4 | 3.774 | 9.884 | 7 | 6.604 | 16.232 |
| Blinatumomab_Arm2_HC | 6 | 5.042 | 11.635 | 4 | 3.361 | 11.451 |
| Blinatumomab_Arm2_LC | 5 | 4.545 | 18.196 | 6 | 5.455 | 16.426 |
| Amivantamab_Arm1_HC | 17 | 3.736 | 51.922 | 15 | 3.297 | 48.186 |
| Amivantamab_Arm1_LC | 8 | 3.738 | 21.417 | 7 | 3.271 | 21.017 |
| Amivantamab_Arm2_HC | 15 | 3.341 | 48.192 | 15 | 3.341 | 54.071 |
| Amivantamab_Arm2_LC | 6 | 2.804 | 24.065 | 6 | 2.804 | 23.665 |
| Faricimab_Arm1_HC | 16 | 3.456 | 43.338 | 16 | 3.456 | 43.142 |
| Faricimab_Arm1_LC | 6 | 2.804 | 23.716 | 6 | 2.804 | 23.716 |
| Faricimab_Arm2_HC | 16 | 3.532 | 52.157 | 17 | 3.753 | 48.415 |
| Faricimab_Arm2_LC | 11 | 5.164 | 23.779 | 9 | 4.225 | 20.457 |
| Mosunetuzumab_Arm1_HC | 17 | 3.786 | 49.435 | 16 | 3.563 | 53.803 |
| Mosunetuzumab_Arm1_LC | 8 | 3.653 | 24.200 | 7 | 3.196 | 26.671 |
| Mosunetuzumab_Arm2_HC | 15 | 3.319 | 57.368 | 16 | 3.540 | 52.206 |
| Mosunetuzumab_Arm2_LC | 6 | 2.817 | 19.772 | 7 | 3.286 | 23.765 |
| Teclistamab_Arm1_HC | 17 | 3.795 | 56.025 | 16 | 3.571 | 51.279 |
| Teclistamab_Arm1_LC | 10 | 4.673 | 20.226 | 9 | 4.206 | 17.086 |
| Teclistamab_Arm2_HC | 15 | 3.319 | 50.733 | 14 | 3.097 | 47.244 |
| Teclistamab_Arm2_LC | 8 | 3.721 | 24.309 | 8 | 3.721 | 25.671 |
| Epcoritamab_Arm1_HC | 16 | 3.524 | 50.871 | 15 | 3.304 | 49.693 |
| Epcoritamab_Arm1_LC | 8 | 3.721 | 26.304 | 8 | 3.721 | 26.348 |
| Epcoritamab_Arm2_HC | 16 | 3.548 | 47.763 | 16 | 3.548 | 46.018 |
| Epcoritamab_Arm2_LC | 7 | 3.271 | 20.023 | 6 | 2.804 | 18.451 |
| Talquetamab_Arm1_HC | 15 | 3.326 | 50.557 | 14 | 3.104 | 47.068 |
| Talquetamab_Arm1_LC | 8 | 3.721 | 24.309 | 8 | 3.721 | 25.671 |
| Talquetamab_Arm2_HC | 15 | 3.371 | 47.960 | 16 | 3.596 | 50.025 |
| Talquetamab_Arm2_LC | 6 | 2.804 | 18.391 | 6 | 2.804 | 18.315 |
| Linvoseltamab_Arm1_HC | 14 | 3.104 | 48.280 | 14 | 3.104 | 46.922 |
| Linvoseltamab_Arm1_LC | 7 | 3.256 | 20.955 | 7 | 3.256 | 20.737 |
| Linvoseltamab_Arm2_HC | 14 | 3.111 | 45.993 | 14 | 3.111 | 44.115 |
| Linvoseltamab_Arm2_LC | 7 | 3.256 | 20.955 | 7 | 3.256 | 20.737 |
| Elranatamab_Arm1_HC | 14 | 3.132 | 48.862 | 15 | 3.356 | 51.250 |
| Elranatamab_Arm1_LC | 8 | 3.653 | 21.259 | 7 | 3.196 | 24.511 |
| Elranatamab_Arm2_HC | 15 | 3.401 | 50.183 | 16 | 3.628 | 49.879 |
| Elranatamab_Arm2_LC | 7 | 3.256 | 20.501 | 7 | 3.256 | 17.425 |

*nHS = Number of Hot Spots; NnHS = Normalized nHS per 100 residues; AAT = Area of profile Above Threshold. All values produced by Aggrescan (Conchillo-Solé et al., 2007).*

### Table S2. Full TAP dataset — wild-type vs PTIm™-mAb optimized, all paired arms

| **Arm** | **WT CDR** | **WT PSH** | **WT PPC** | **WT PNC** | **WT SFvCSP** | **OPT CDR** | **OPT PSH** | **OPT PPC** | **OPT PNC** | **OPT SFvCSP** |
| --- | --- | --- | --- | --- | --- | --- | --- | --- | --- | --- |
| Blinatumomab_Arm1 | 46 | 139.0259 | 1.9952 | 0.1145 | 8.4 | 47 | 149.0163 | 0.6885 | 0.3043 | -2.0 |
| Blinatumomab_Arm2 | 49 | 118.8103 | 1.4192 | 1.2516 | 8.13 | 49 | 161.0832 | 1.1062 | 2.3475 | 1.1 |
| Amivantamab_Arm1 | 52 | 153.8806 | 0.0795 | 1.3919 | 0.0 | 47 | 128.9158 | 0.0 | 1.1217 | 0.0 |
| Amivantamab_Arm2 | 46 | 117.0281 | 0.0 | 0.0 | 3.41 | 46 | 114.257 | 0.0 | 0.0 | 9.0 |
| Faricimab_Arm1 | 56 | 143.4099 | 0.0853 | 0.0 | 4.4 | 56 | 139.2502 | 0.0852 | 0.0 | 4.4 |
| Faricimab_Arm2 | 52 | 138.909 | 0.0114 | 1.3583 | -0.38 | — | — | — | — | — |
| Mosunetuzumab_Arm1 | 51 | 136.2322 | 1.3391 | 0.1574 | 0.0 | 45 | 128.3042 | 0.0568 | 0.1106 | -3.0 |
| Mosunetuzumab_Arm2 | 48 | 116.6835 | 0.0 | 0.0 | 6.0 | 47 | 108.3864 | 0.0 | 0.0 | 2.0 |
| Teclistamab_Arm1 | 50 | 155.2379 | 0.332 | 1.5434 | -3.89 | 50 | 173.7491 | 8.0279 | 1.4258 | -1.0 |
| Teclistamab_Arm2 | — | — | — | — | — | 52 | 158.4317 | 1.5056 | 0.0 | 0.0 |
| Epcoritamab_Arm1 | 55 | 154.4802 | 1.3399 | 0.0 | 0.0 | 50 | 150.5429 | 1.4312 | 0.0 | 0.1 |
| Epcoritamab_Arm2 | 49 | 132.6275 | 0.0 | 0.0446 | -0.9 | 47 | 131.9073 | 0.0 | 0.1054 | -7.6 |
| Talquetamab_Arm1 | — | — | — | — | — | 52 | 156.7761 | 3.5033 | 0.0 | 0.0 |
| Talquetamab_Arm2 | 45 | 101.0231 | 0.0154 | 0.0 | 15.3 | 47 | 111.2287 | 0.0434 | 0.0 | 20.4 |
| Linvoseltamab_Arm1 | 53 | 157.8034 | 0.1649 | 0.466 | -3.0 | 47 | 130.8496 | 0.1128 | 0.1755 | -6.0 |
| Linvoseltamab_Arm2 | 52 | 142.7591 | 0.806 | 1.2885 | -2.4 | 47 | 132.1593 | 0.0 | 1.3917 | -3.0 |
| Elranatamab_Arm1 | 53 | 144.7301 | 1.7981 | 0.318 | 8.2 | 49 | 145.6310 | 0.5799 | 0.1083 | -5.1 |
| Zenocutuzumab | 51 | 127.3201 | 0.1191 | 0.0 | 12.5 | 47 | 127.8476 | 0.0 | 0.0 | 6.0 |
| Emicizumab_Arm1 | 50 | 121.48 | 2.0118 | 0.4202 | 3.0 | 47 | 111.44 | 2.9829 | 0.2605 | 5.0 |
| Emicizumab_Arm2 | 46 | 112.78 | 4.0449 | 0.1091 | -1.0 | 46 | 117.30 | 4.0298 | 0.1262 | 2.0 |

*"—" = TAP submission error. CDR = Total CDR length; PSH = CDR-vicinity Patches of Surface Hydrophobicity; PPC = CDR-vicinity Patches of Positive Charge; PNC = CDR-vicinity Patches of Negative Charge; SFvCSP = Structural Fv Charge Symmetry Parameter.*

### Table S3. Cost and time assumptions underlying the savings figures in Section 3.5

The per-category cost and time assumptions tabulated below are SANSHI internal industry-benchmark estimates derived from typical CRO pricing for therapeutic-antibody discovery campaigns and from published average timelines for the corresponding activities. They are presented for transparency: prospective licensees and readers can substitute their own internal cost structures when applying the framework to their own programs. We do not claim that these figures derive from any single peer-reviewed source.

| **Cost category** | **Time assumption** | **Cost assumption (USD)** | **Rationale** |
| --- | --- | --- | --- |
| CRO humanization (1 round) | 6 weeks per round | $75,000 per round | Construct synthesis + transient expression + binding assay at typical CDMO pricing. |
| Deimmunization design (verification) | 1 month | $75,000 | One verification round for the deimmunized construct. |
| Sequence-liability targeted assessment | 0.5–1 month | $30,000 | Directed LC-MS forced-degradation study at known liability positions, scoped down from a full-sequence scan. |
| Aggregation-prone-region engineering | 1.5 months | $50,000 | Reduced scope of formulation screening when aggregation is addressed at the sequence level; based on roughly half of a full DOE formulation screen ($100K typical). |
| Developability early flag | 2 months | $150,000 | Cost of late-stage re-engineering (≈2 humanization rounds) avoided when low-developability candidates are flagged before CMC commitment. |
| Architecture selection (per format) | 1 month per format | $50,000 per format | Cost of constructing, expressing, and characterizing one bispecific format; PTIm™ computationally compares formats without requiring the physical build. |
| SPR/BLI binding-characterization campaign | 0.5 month | $15,000 | Per antibody-antigen pair, at typical CDMO pricing. |

*All cost figures are nominal SANSHI internal industry-benchmark estimates as of 2025. They reflect typical CRO and CDMO pricing in the US and Western European market; figures in other regional markets may differ. These assumptions form the basis of the per-project totals reported in Section 3.5 and visualized in Figure 5. Customers are advised to substitute their own internal cost structures when applying the framework to their own programs.*
